# A simple and effective method to isolate germ nuclei from *C. elegans* for genomic assays

**DOI:** 10.1101/371351

**Authors:** Mei Han, Guifeng Wei, Catherine E. McManus, Valerie Reinke

## Abstract

The wide variety of specialized permissive and repressive mechanisms by which germ cells regulate developmental gene expression are not well understood genome-wide. Isolation of germ cells with high integrity and purity from living animals is necessary to address these open questions, but no straightforward methods are currently available. Here we present an experimental paradigm that permits the isolation of the nuclei from *C. elegans* germ cells at quantities sufficient for genomic analyses. We demonstrate that these nuclei represent a very pure population and are suitable for both chromatin immunoprecipitation (ChIP-seq) and transcriptome (RNA-seq) analyses. The method does not require specialized transgenic strains or growth conditions and can be readily adopted by other researchers with minimal troubleshooting. This new capacity removes a major barrier in the field to dissect gene expression mechanisms in the germ line of *C. elegans*. Consequent discoveries using this technology will be relevant to conserved regulatory mechanisms across species.

## Introduction

Establishing tissue-specific gene expression programs during development requires dynamic, highly coordinated gene regulation, often over extended genomic regions. The *C. elegans* germ line is an ideal microcosm to explore complex gene expression regulatory mechanisms. These germ cells deploy diverse, tightly controlled gene regulatory programs to drive proliferation, meiosis and gamete differentiation, yet retain the ability to reactivate totipotency in the zygote [1]. They must therefore repress somatic gene expression, which could lead to inappropriate or premature differentiation [2]. Indeed, ectopic activation of somatic programs readily transforms germ cells to neurons, intestine, and muscle [3, 4]. Germ cells exhibit long-range regulation as well, across multi-megabase-long piRNA gene clusters [5] and over the entire X chromosome [6]. All of these complex events must be precisely coordinated to permit the production of hundreds of viable embryos in each hermaphrodite in just a few short days of reproductive capacity.

Chromatin-based, post-transcriptional, and small RNA mechanisms play a central role in modulating transcript and protein abundance in the germ line [7]. For example, the conserved Rb/E2F regulatory complex is critical for establishing distinct germline and somatic gene expression programs [8-11]. Additionally, germ cells are transformed to somatic cells in vivo either by disrupting chromatin regulation via forced expression of a somatic transcription factor concomitant with loss of chromatin factor LIN-53 [4], or by disrupting post-transcriptional regulation through loss of mRNA-binding proteins MEX-3 and GLD-1 [12] or loss of germ granules [13, 14]. Distinct small RNA pathways selectively target transcripts either for degradation or protection in the cytoplasm, and ultimately alter chromatin state as well. Disrupting the feedback from cytoplasm to nucleus causes germ cells to gradually lose their identity over multiple generations [15].

To fully investigate these regulatory mechanisms, genome-scale assays are necessary. However, in many species, it is difficult to isolate sufficient germ cells at key developmental times due to their relative scarcity and sequestration within various somatic niches. In *C. elegans*, germ cells are prominent in number and location relative to somatic cells, but to date, no methods have been developed to purify them from living animals, in part because these cells share cytoplasm via cellular bridges and therefore exist in a syncytium. Moreover, thousands of animals are required to provide sufficient material for most assays, eliminating the option of gonad dissection. Although proteins can be epitope-tagged and expressed specifically in the germ line for tissue-specific ChIP-seq [9], many other applications such as histone modification profiling and chromatin organization assays require pure chromatin preparations. Histone modifications and chromatin states have therefore been measured only in whole animal or embryo preparations, which mix somatic and germ cell populations and complicate interpretation of the data. Here we report a simple method that circumvents this limitation and produces populations of germ nuclei at ~90% purity, with yields sufficient for biochemical and genomic analyses. We show that these isolated germ nuclei (IGN) can be used for both RNA-seq and histone modification ChIP-seq, and exhibit expected patterns of germline gene regulation. This procedure is straightforward and easily adaptable to any worm strain with appreciable numbers of germ cells, and these nuclei can be useful for additional genomic assays, including chromatin capture conformation, nucleosome accessibility, and small RNA-seq, among others.

## Results

### A highly efficient method to isolate germ nuclei from *C.elegans* adults

We have developed a novel, simple procedure to isolate germ nuclei at a scale that permits biochemical analyses (Figure 1A). Based on a previous report in which intestine nuclei could be isolated from whole animals [16], we determined conditions in which germ nuclei are preferentially released from the syncytial gonad by gentle Dounce homogenization and vortexing of whole adult animals, followed by filtration to remove cellular (somatic) debris (see Materials and Methods). The method takes approximately three hours from worm harvest to nuclear pellet and results in 1.5-3×10^7^ nuclei from ~1×10^6^ worms. Isolated nuclei show relatively uniform size and intact nuclear structure based on DAPI staining (Figure 1B).

**Figure 1.**
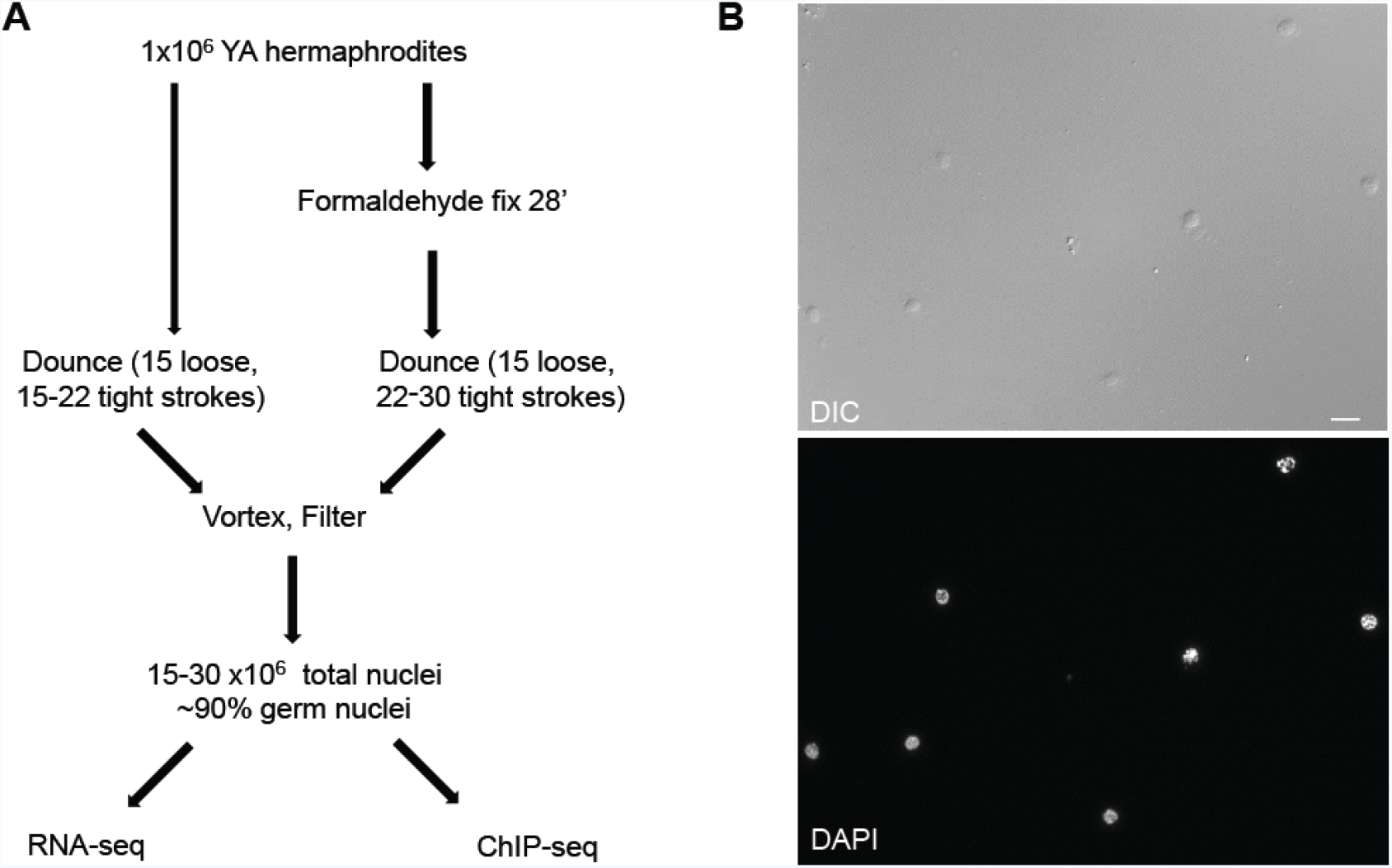
A simple method to isolate germ nuclei from *C. elegans*. (A) Schematic representation of the method to isolate germ nuclei from *C. elegans.* Approximately 1×10^6^ young adult hermaphrodites were collected for nuclei isolation for each experiment. For RNA-seq, worms were homogenized with 15 loose and 15-22 tight Dounce strokes after collection. For ChIP-seq, worms were fixed with 2% formaldehyde for 28 minutes before homogenization with 15 loose and 22-30 tight Dounce strokes (see Materials and Methods). A typical yield is 15-30 ×10^6^ nuclei from 1×10^6^ young adult hermaphrodites. (B) Representative image of isolated total nuclei from young adult worms stained with DAPI. Scale bar, 10 μm.

To calculate the percentage of isolated nuclei that are from germ cells, we performed the isolation procedure on a transgenic strain that expresses OEF-1::GFP. OEF-1 is a novel germline factor present specifically in mitotic and pachytene nuclei that disappears abruptly at the onset of oogenesis [17] (Figure 2A). Immunostaining of isolated nuclei from this strain with anti-GFP indicates that 91% are positive for OEF-1 (Figure 2B and Table S1). We also used a second germline-specific transgenic strain, AZ212, that expresses GFP::H2B specifically in the germ line under the control of the *pie-1* promoter [18] (Figure S1). In this strain, GFP::H2B is expressed in the nuclei of germ cells at all developmental stages, including oocytes. However, we found a staining frequency (89%) in isolated nuclei similar to that of OEF-1::GFP, which suggests that most of the nuclei obtained are from the distal and/or medial gonad. Indeed, many of the isolated nuclei appear to be in the pachytene stage based on chromosome morphology visible by DAPI staining (Figure 2B and S1B). Many of the unstained nuclei exhibit extremely condensed DNA, and we suspect that they represent sperm that are released during the disruption protocol and pass through the size filters (Figure S2). Thus, the actual percentage of somatic nuclei present in the population is likely much less than 10%.

**Figure 2.**
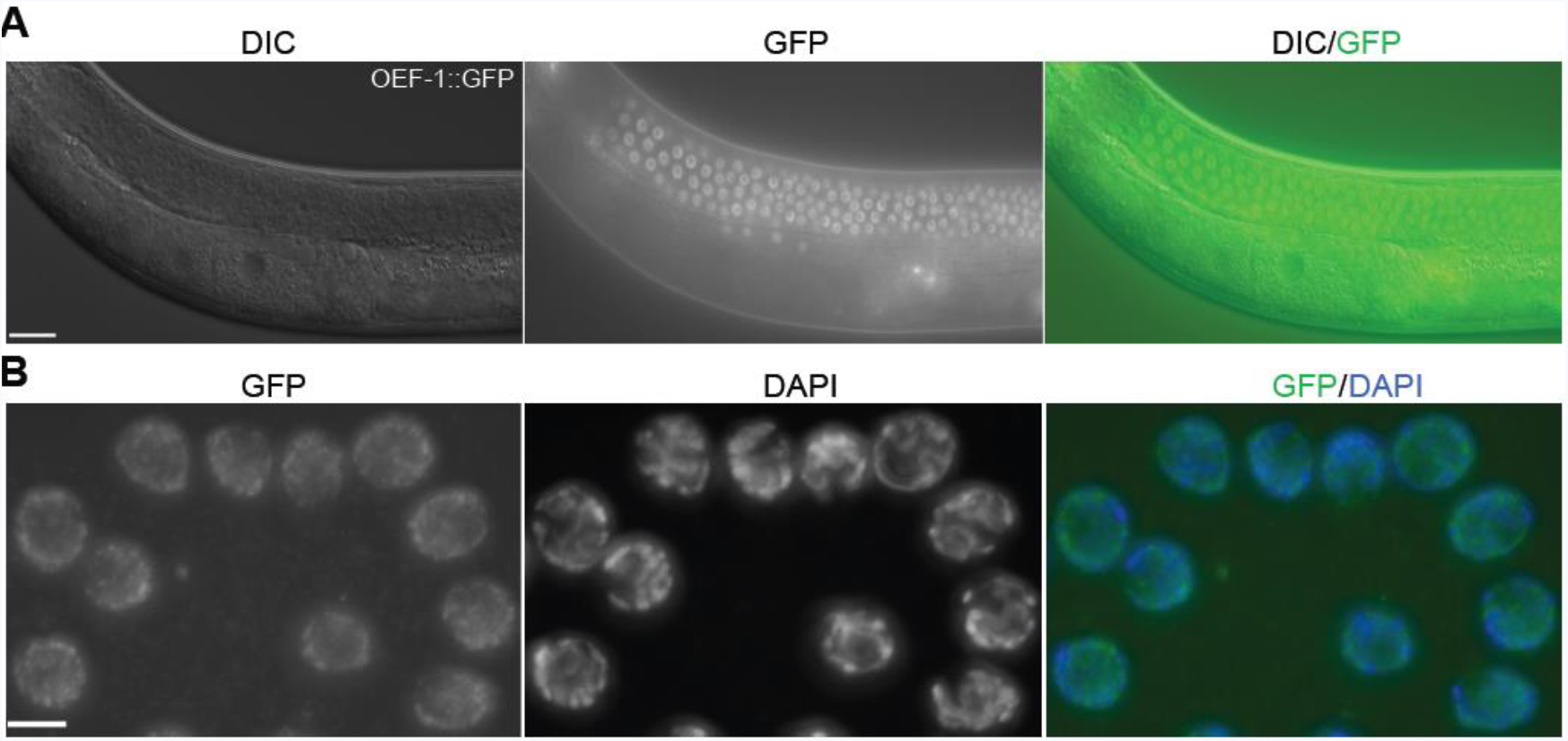
Quantification of isolated germline nuclei. (A) Young adult OEF-1::GFP transgenic worms with germ line expression. OEF-1 is detected specifically on autosomes in mitotic and pachytene nuclei and disappears at the onset of oogenesis [17]. (B) Isolated nuclei from OEF-1::GFP young adult worms were immunostained with GFP (green) and stained with DAPI (blue). Nuclei stained with both GFP and DAPI were designated germline nuclei (91.02% of total nuclei, n=2127). Two independent biological replicates were performed. Scale bars, 20 μm in (A), 5 μm in (B).

We have performed this isolation protocol on both unfixed and fixed animals, depending on the downstream application. Fixation slightly increases the number of Dounce strokes necessary to break open the animals but does not otherwise impair the procedure (Figure 1; Materials and Methods). One of the main advantages of this approach is that it does not require any specialized transgenic system or subsequent affinity purification, and it is applicable to any transgenic or mutant strain with appreciable numbers of germ nuclei. In sum, this method is rapid, simple, reproducible, and adaptable. We use the abbreviation “IGN” to refer to these isolated germ nuclei. As described below, we have performed IGN-RNA-seq as well as IGN-ChIP-seq for a histone modification, demonstrating that the nuclei are amenable to a wide variety of genomic assays.

### IGN expression analysis

We first performed total RNA-seq with ribosomal RNA depletion on IGN from wild type (N2) young adults. To permit direct comparison to somatic gene expression, we also performed whole-animal RNA-seq on *glp-1(q224)* young adults, which have a temperature-sensitive mutation in the GLP-1/Notch receptor. At the restrictive temperature, *glp-1(q224)* mutants lack all germ cells except for a few mature sperm, and thus represent only somatic tissues (which we abbreviate as SOM throughout the manuscript) [19]. Two independent RNA-seq experiments were analyzed for each genotype using HISAT2 [20] and Cuffdiff [21]. On average, 24 million paired-end sequenced reads were mapped to the *C. elegans* genome (ce10) per sample. Transcript abundance is reported as Fragments Per Kilobase of transcript per Million mapped reads (FPKM). We observed many obvious differences between IGN and SOM profiles shown in genome browser views (representative images in Figure S3). We thus identified differentially expressed genes between IGN and SOM with a q value less than 0.05, which equals 1.36-fold or greater difference in expression. These analyses identified 5075 genes with IGN-enriched expression and 3965 genes with SOM-enriched expression (Figure 3A and 3B; File S1). We examined the Gene Ontology (GO) biological process terms for IGN-enriched transcripts. Strikingly, most terms are related to the mitotic or meiotic cell cycle and gamete generation, indicating that genes with increased expression in IGN relative to SOM are highly associated with germline-associated functions (Figure 3C).

**Figure 3.**
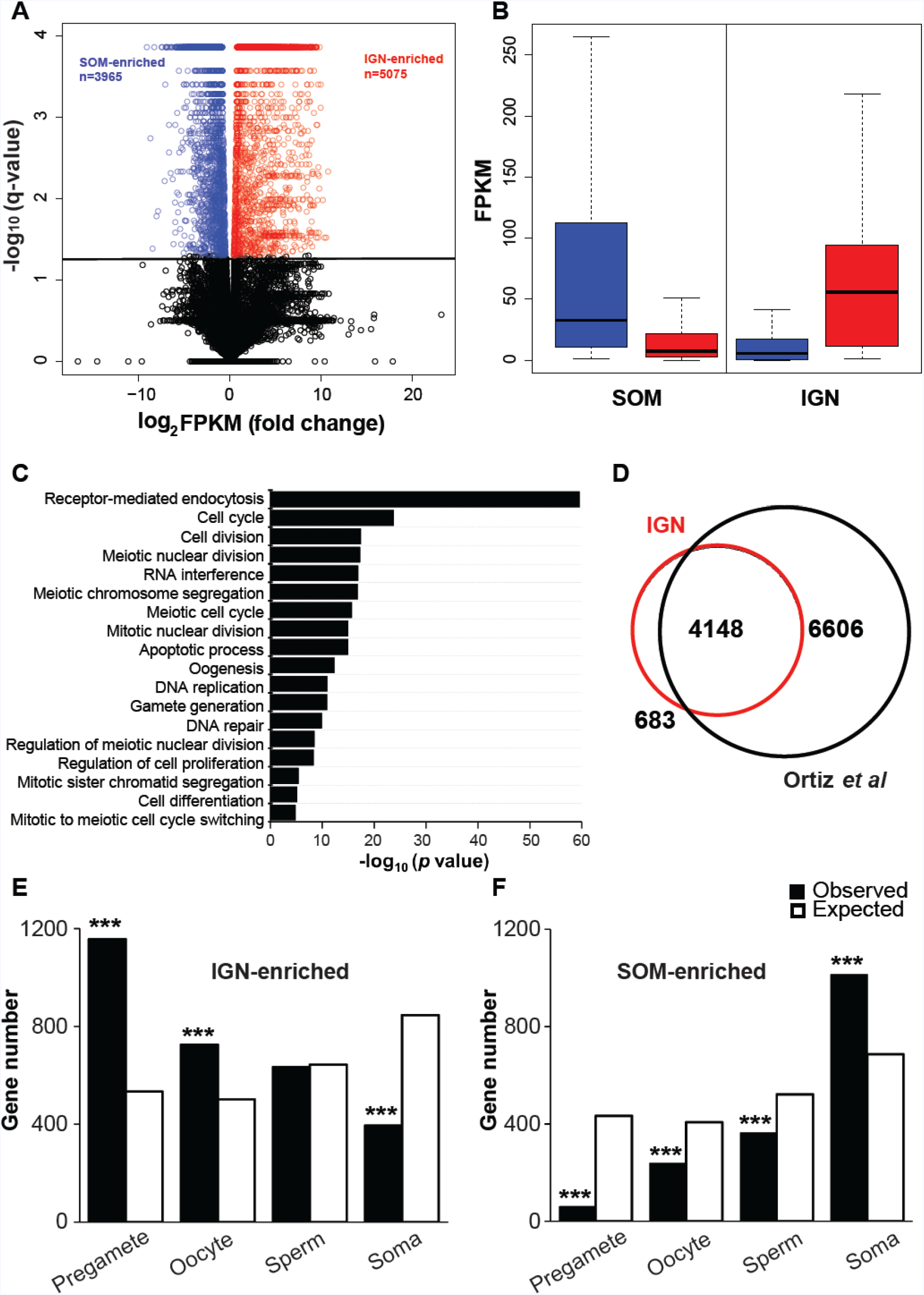
Expression profiling of *C. elegans* isolated germline nuclei. RNA from two independent replicates of wild type IGN and from SOM (*glp-1(q224)* young adults) was analyzed for expression profiling. (A) Volcano plot showing −log10 of q-value against log2 of fold change for each gene. The number of genes that were significantly up-regulated in SOM or IGN are indicated. Black line marks the significance cutoff of q = 0.05 (Y axis). Blue circles indicate SOM-enriched transcripts and red circles indicate IGN-enriched transcripts. (B) A boxplot displaying the overall abundance and distribution of gene expression levels (FPKM) for 3965 SOM-enriched transcripts (blue) and 5075 IGN-enriched transcripts (red) in either IGN RNA-seq or SOM RNA-seq. Each box indicates the median and interquartile range of FPKM level. (C) The most significant Gene Ontology Biological Process terms of the 5075 IGN-enriched transcripts. (D) Overlap of 4831 IGN-enriched coding transcripts with previously-identified germline-expressed transcripts in dissected gonad [22]. (E-F) Bar graphs indicating expected and observed number of genes (Y axis) in different gene categories [23] (X axis) for IGN-enriched transcripts (E) and SOM-enriched transcripts (F). Asterisks indicate significantly more genes than expected (hypergeometric test, p-value<1×10^-5^ [^**^], p-value<1×10^−10^ [^***^]).

To determine how well the IGN-enriched dataset reflects known germline-expressed or germline-enriched gene expression profiles, we made several comparisons with published datasets. A total of 10,754 genes were identified as expressed in dissected gonads from hermaphrodites [22]. Of the 5075 genes with IGN-enriched expression, 4831 are coding and 244 are non-coding genes. We found that 4148 out of the 4831 are present in the dissected gonad dataset (Figure 3D; File S2). Of the 683 genes present exclusively in the IGN dataset, many are encoded by genes with known germline function, such as *hal-2*, *rec-1*, *mes-6*, *snpc-4*, *vbh-1*, etc. Many others have quite low expression and might be a consequence of different experimental design (e.g. poly-A purification vs ribo-depletion strategies) or different cutoffs in the analysis. Conversely, a large number (6606) of genes are present exclusively in the dissected gonad dataset. This dataset by definition is more encompassing, as it represents a comprehensive set of germline-expressed genes, whereas the IGN-enriched dataset excludes genes expressed at similar or higher levels in the soma. Consistent with this possibility, GO analysis indicates that the 6606 genes represented exclusively in the dissected gonad dataset mainly contribute to fundamental cellular processes, including oxidation-reduction, lipid metabolism, transport, proteolysis etc., which occur in most or all cell types and are not expected to be especially enriched in germ cells (Figure S4).

To further validate the IGN-enriched and SOM-enriched datasets, we also compared them to genes previously placed in distinct expression categories: pre-gametic germ cells, oocytes, sperm, and the soma [23]. Genes in the IGN-enriched dataset are over-represented among the pre-gamete and oocyte but not the sperm categories, and under-represented in the soma category (Figure 3E; File S2). Conversely, genes in the SOM-enriched dataset are over-represented in the soma category and under-represented in the germline-related categories (Figure 3F and File S2). Altogether, these results indicate that IGN samples are suitable for RNA-seq to examine nuclear germline-specific gene expression profiles.

### Chromatin modification profiling of germ nuclei

We next tested whether IGN could be used to profile histone modifications specifically in germ nuclei. We selected H3K27ac as a good candidate for ChIP because it is highly associated with active enhancers/promoters [24-26]. In addition to IGN, we also isolated chromatin from *glp-1* mutant young adults to represent somatic tissues. We first used ChIP-qPCR to test a handful of loci. Two correspond to the upstream regulatory regions of genes with known germline-specific expression, *him-3* [27] and *oef-1* [17]. Three were selected from existing H3K27ac ChIP-chip data from the ENCODE project at the L3 stage of development (GSM624432 and GSM624433) [28] to serve as positive (*C37H5.15*) and negative (*elt-2, myo-3*) controls. Both *him-3* and *oef-1* upstream regions showed significant enrichment (10-30x) for H3K27ac in IGN relative to SOM (Figure S5).

We therefore performed H3K27ac ChIP-seq on IGN and SOM chromatin, and called peaks using MACS2 (see Materials and Methods). Immunostaining of dissected gonads has indicated that such activating histone modifications are primarily associated with autosomes and depleted from the X chromosome in germ cells, where the X is largely silenced [29, 30]. Consistent with this expectation, fewer H3K27ac peaks than expected are observed on the X in the IGN-ChIP-seq dataset. By contrast, the X had more peaks than expected in the SOM dataset (Figure 4A and 4B). Certain autosomes had significant variations from expected values of peaks, but for a given autosome, the trend was the same for both IGN and SOM.

**Figure 4.**
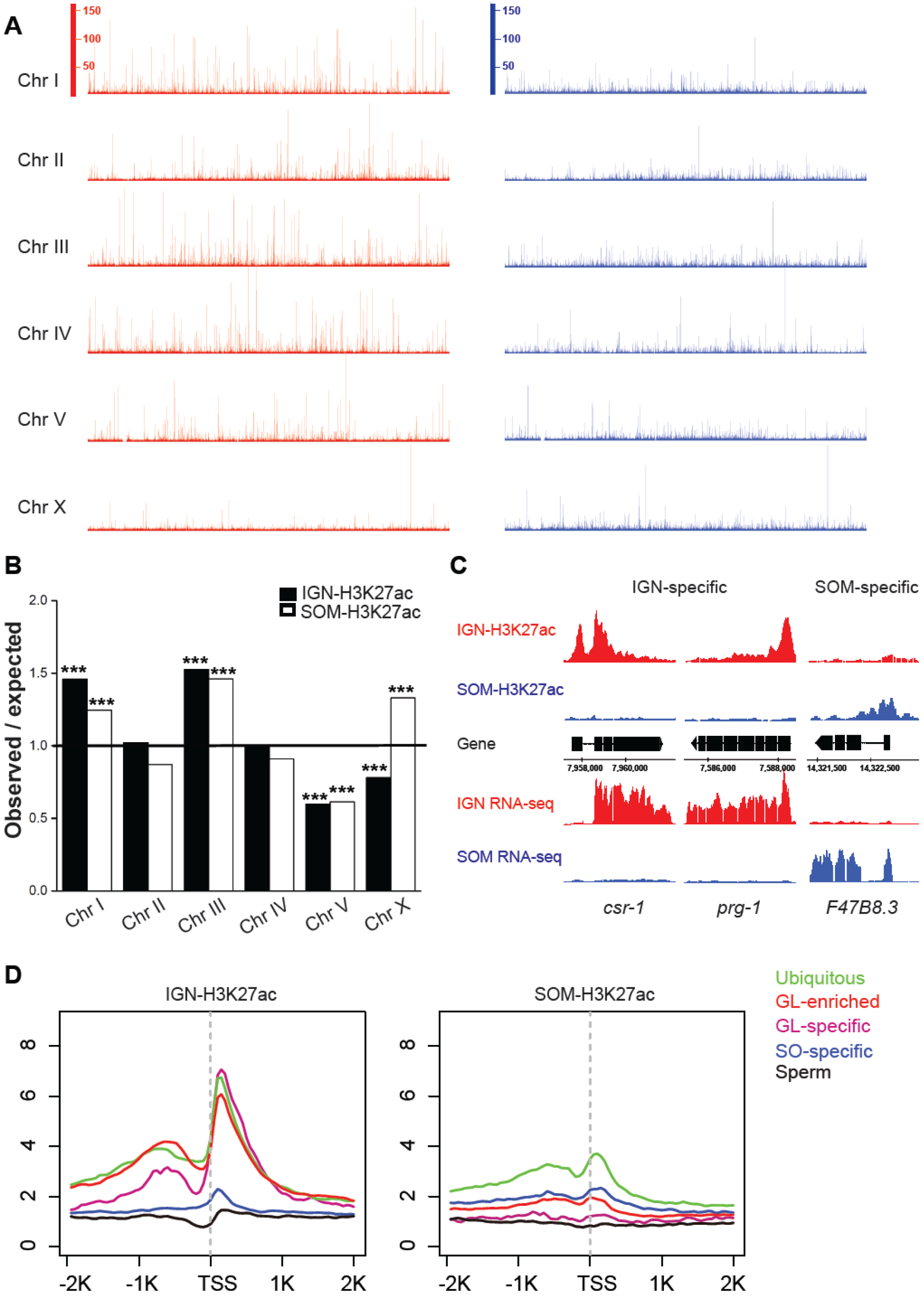
H3K27ac exhibits tissue-specific profiles in IGN and SOM. (A) Genome view of normalized ChIP-seq reads for IGN-H3K27ac ChIP-seq (left panel) and SOM-H3K27ac ChIP-seq (right panel) over all six chromosomes. (B) Distribution of H3K27ac modified coding targets across chromosomes relative to genomic length for IGN-H3K27ac ChIP-seq and SOM-H3K27ac ChIP-seq datasets. Statistically significant deviations from the expected value of 1 (indicated by black line) are indicated by asterisks (Pearson’s chi-square test, p-value<1×10^−5^ [^**^], p-value<1×10^−10^ [^***^]). (C) Example ChIP-seq and RNA-seq tracks across the IGN-specific genes *csr-1* and *prg-1* and SOM-specific gene *F47B8.3*. IGN signals are indicated in red and SOM signals are indicated in blue. (D) Metagene analysis of the distribution of average ChIP-seq signal for indicated gene categories [13] for IGN and SOM H3K27ac profiles.

Across all chromosomes, tissue-specific H3K27ac ChIP-seq signals were easily distinguished between IGN and SOM in the genome browser (Figure S6). Many genes known to have germline-specific expression, such as *csr-1* and *prg-1*, exhibit high levels of H3K27ac at or near their transcription start sites in IGN and minimal to no signal in SOM, while many genes such as *F47B8.3* exhibit the converse pattern (Figure 4C). We identified 15148 significant peaks for the IGN dataset and 8667 peaks for the SOM dataset. Of these, 5059 are shared (overlapping by at least 1 nt), leaving over 10,000 peaks unique to IGN. We next assigned H3K27ac peaks to neighboring candidate genes genome-wide (File S3; Materials and Methods). Target genes with common peaks between IGN and SOM are largely categorized as having basic metabolic functions, including translation, protein folding, and tricarboxylic acid cycle, while genes marked by IGN-specific H3K27ac are linked to germline functions, including cell cycle, cell division, mitotic/meiotic nuclear division and receptor-mediated endocytosis (Figure S7).

To compare the distribution of H3K27ac marks around the transcription start sites (TSS) for different gene categories in IGN and SOM, we performed metagene analyses for genes assigned to one of five previously published expression categories: ubiquitous, germline-specific, germline-enriched, soma-specific, and sperm [13] (Figure 4D). Genes classified as ubiquitous showed abundant H3K27ac enrichment in IGN, suggesting an overall active chromatin state in germ cells. Similarly, the germline-enriched and germline-specific genes displayed higher H3K27ac levels than soma-specific genes in IGN. By contrast, soma-specific genes exhibited higher H3K27ac levels in SOM, as expected. The lack of the H3K27ac modification for genes in the sperm category is consistent with the fact that such genes, which are normally expressed only at L4 [31], are not actively transcribed in the adult IGN or SOM.

We next analyzed the relationship between H3K27ac levels and transcript abundance for IGN and SOM samples. Genes with IGN-enriched expression exhibit higher H3K27ac levels in IGN-H3K27ac ChIP-seq compared to genes with SOM-enriched expression (Figure 5A). Conversely, genes with SOM-enriched expression exhibit higher H3K27ac in the SOM-H3K27ac ChIP-seq dataset than genes with IGN-enriched expression. Notably, the overall levels of H3K27ac in IGN are strikingly higher for IGN-enriched genes than in SOM for SOM-enriched genes (Figure 5A and 5B). Indeed, the strength of the H3K27ac signal correlated with transcript abundance better in IGN than SOM (Figure 5C). Genes with high expression (FPKM ≥100) and high H3K27ac peak enrichment (≥11) included many genes with “housekeeping” functions that are expected to be highly expressed in germ cells, such as those encoding ribosomal proteins, as well as genes with known functions in the germ line (File S3; Figure S8). In SOM samples, even among highly expressed genes (≥100 FPKM), few reached the same threshold of H3K27ac peak enrichment of ≥11. These results suggest that in germ cells, H3K27ac abundantly marks active genes that are required for germ cell cycle and germline development, regardless of whether these genes are broadly or more specifically expressed. By contrast, H3K27ac enrichment is not particularly correlated with transcript abundance or particular gene functions in somatic tissues at the adult stage.

**Figure 5.**
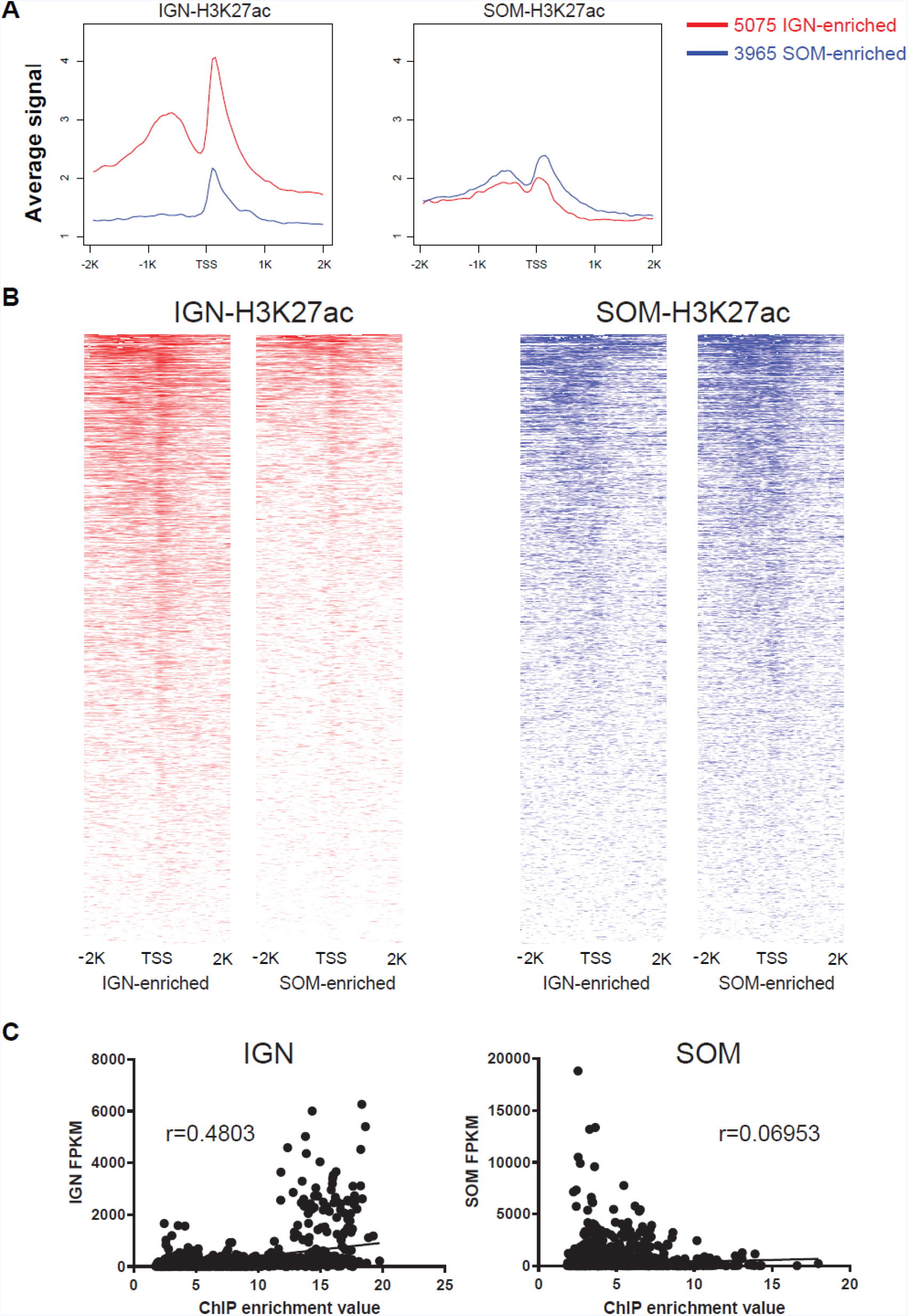
H3K27ac levels correlate with highly expressed genes in IGN but not SOM. (A) Metagene analysis shows the average signal distribution of IGN-H3K27ac ChIP-seq (left panel) and SOM-H3K27ac ChIP-seq (right panel) for 5075 IGN-enriched transcripts (red line) and 3965 SOM-enriched transcripts (blue line). (B) Heat map of IGN-H3K27ac ChIP-seq (red) and SOM-H3K27ac ChIP-seq (blue) around TSS of 5075 IGN-enriched transcripts and 3965 SOM-enriched transcripts. Read density was extracted from a genomic region ± 2 kb around TSS for each gene. The gradient red-to-white color (IGN-H3K27ac ChIP-seq) or blue-to-white color (SOM-H3K27ac ChIP-seq) indicates high-to-low density in the corresponding region. (C) Tissue-specific correlation between RNA-seq and ChIP-seq data. A unique ChIP enrichment value was chosen for each target gene (see Materials and Methods). Scatter plot for IGN-enriched transcript abundance (FPKM) and IGN-H3K27ac ChIP enrichment is shown in left panel, while the same comparison is shown for SOM in right panel. Pearson r=0.4803 for IGN sample and r=0.06953 for SOM sample.

In sum, this analysis indicates that IGN can be used to isolate chromatin and perform ChIP-seq to identify histone modification profiles that are present specifically in the adult germ line.

## Discussion

Here we present a novel method for isolating germ nuclei (IGN) from *C. elegans* adult hermaphrodites in quantities sufficient for genomic analyses. We demonstrate that IGN preparations contain approximately ~90% germ nuclei under conditions in which animals are fixed prior to Dounce treatment. Moreover, we establish that IGN are suitable for genomic assays such as RNA-seq and histone modification ChIP-seq, and display profiles consistent with prior analyses of germline gene expression. Thus, this technically straightforward and adaptable procedure should greatly simplify future genomic studies of germline gene regulation.

As we established conditions for the isolation procedure, we found that a balance must be reached in the number of Dounce strokes to achieve sufficient breakage and yet preserve the integrity of the nuclei. Thus, most animals remain intact throughout the process, and currently, the yield can vary from 15-30 nuclei per animal, depending on the condition of the Dounce homogenizers. We suspect that certain mutant strains that alter cuticle structure, worm shape, or otherwise affect susceptibility to breakage might further improve efficiency and reduce the amount of effort spent culturing worms.

The RNA-seq data clearly demonstrates that IGN display a strong enrichment for genes known to have germline-enriched expression. One important distinction between the IGN dataset and all other previously defined expression datasets is that IGN presumably have comparatively little contribution from cytoplasmic pools; thus nuclear transcripts are likely over-represented. In the future, this feature should be useful for analysis of nascent transcription and of precursor transcripts, for instance, for profiling changes in hnRNAs in splicing or RNA-processing mutants.

Similarly, the ChIP-seq analysis of H3K27ac levels in IGN clearly demonstrates strong enrichment for genes known to have germline-enriched gene expression. Surprisingly, we found a much better correlation between H3K27ac and gene expression in IGN than in SOM, with overall lower levels of H3K27ac in SOM. This reduced association between H3K27ac and gene expression in the soma could be due in part to the mixing of multiple cell types in the sample. Additionally, these analyses were conducted in adult somatic tissues, which have been developmentally stable for some time. Thus, transcript accumulation might be relatively uncoupled from genes currently marked as transcriptionally active by the presence of H3K27ac in upstream regulatory regions. This analysis shows that even straightforward comparisons under wild type conditions can yield new insights when performed in isolated cell types.

In sum, this newly developed procedure to isolate germ nuclei is simple and robust and should be readily employed and adapted to many different experimental conditions. We therefore expect that it will facilitate many inquiries using genomic-based technologies into multiple aspects of gene expression regulation in the *C. elegans* germ line.

## Materials and Methods

### *C. elegans* strains

Strains were maintained by standard methods unless otherwise indicated [32]. Whole-genome sequenced VC2010 (a substrain of N2) was used as the wild type strain. All worm culture was performed at 20°C, except for *glp-1(q224)*, which was maintained at 15°C and shifted to 25°C after synchronized L1s were hatched to induce sterility.

OP383 *unc-119(tm4063)* III; *wgIs383* [*oef-1*::TY1::EGFP::3x-FLAG + *unc-119*(+)] [33]. AZ212 *unc-119(ed3)*; *ruIs32* [*pie-1*p::GFP::H2B + *unc-119*(+)] III [18]. JK1107 *glp-1(q224)* III [19].

### Isolation of germline nuclei

Worms were grown to starvation on 15-cm NGM plates. Starved worms were washed with M9 and collected in a 15 mL conical tube. Worms were floated in 15 mL M9 for 5 minutes and the upper 6 mL worm solution containing mostly L1s was transferred to a new 15 mL tube. 45K L1s were plated to each peptone enriched plate. Worms were grown to gravid on peptone enriched plates, bleached, and hatched overnight in M9 for 16-24 hours. 50K L1s were plated to each enriched plate and grown until the young adult stage. Animal preparations for nuclei isolation were prepared at different scales as follows: worms from eighteen enriched plates (~1 million) were used per ChIP-qPCR or ChIP-seq experiment, and worms from six enriched plates (~3×10^5^) were used for RNA-seq. Young adult animals were harvested at various times after plating synchronized L1s (VC2010: 54-56 hours, OP383: 67-69 hours, AZ212: 68-69 hours), when most of the animals had 4-10 embryos. Worms from every six plates (~3×10^5^) were collected into one 50 mL conical tube and spun at 3100 rpm for 2 minutes and then washed 3× in M9.

For ChIP-qPCR and ChIP-seq, worms were crosslinked in 50 mL 2% formaldehyde for 30 minutes in three 50 mL conical tubes at room temperature [34]. Formaldehyde was quenched by 1 M Tris (pH 7.5) wash. Worms were then washed two more times with M9. Worms were then washed in the same tubes with 10 mL of prechilled Nuclei Purification Buffer (NPB; 50 mM HEPES pH=7.5, 40 mL NaCl, 90 mM KCl, 2 mM EDTA, 0.5 mM EGTA, 0.2 mM DTT, 0.5 mM PMSF, 0.5 mM spermidine, 0.25 mM spermine, 0.1% tween 20, and cOmplete proteinase inhibitor cocktail (Roche) – 1 tablet per 25 mL NPB) [35, 36]. Worms from every six peptone enriched plates were resuspended with prechilled NPB to a final volume of 6 mL and subsequently transferred to a prechilled 7 mL glass Dounce homogenizer (Wheaton, Clearance: 0.05 +/- 0.025mm). All subsequent steps were performed at 4°C or on ice. One Dounce was used per set of 6 plates. A total of 15 loose and 22-30 tight strokes with a quarter turn after each stroke were performed to homogenize worms. After every 15 Dounce strokes, the sample was held for five minutes on ice. The optimal number of tight strokes depends on worm stage, worm genotype and the condition of the Dounce homogenizer. A fraction of broken worms between 8-15% after the last tight stoke normally related to good release of germline nuclei. NPB was then added to a final volume of 10 mL for worms from every 6 plates. The combined 30 mL worm solution from 3 Dounce sets were transferred to a prechilled 50 mL conical tube and vortexed on medium-high speed for 30 seconds, followed by 5 minutes on ice. The vortex and ice incubation steps are repeated one time to release more nuclei. The solution was passed through six 40 μm cell strainers (Fisherbrand) and six 20 μm cell strainers (pluriSelect) to remove worm debris. Two more 20 μm cell strainers were used for the filtrate to further remove worm debris. Isolated nuclei were collected by centrifugation at 3100 rpm for 6 minutes at 4°C. The supernatant was removed and the nuclei were resuspended with 1 mL NPB. The nuclei were then transferred to a nonstick 1.5 mL tube (Ambion). A 5 μL aliquot of nuclei were stained with DAPI and counted with a hemacytometer (Hausser Scientific). Normally 15-30 million nuclei can be isolated from around 1 million young adult animals. The rest of nuclei were pelleted at 4000 rpm for 5 minutes at 4°C. Most of the supernatant was removed so that ~20 μL NPB was left. The nuclei were gently pipetted to mix and then flash frozen in liquid nitrogen and stored at −80°C until sonication.

For RNA-seq, worms from six peptone enriched plates were incubated for 30–45 minutes in M9 buffer with shaking after the third M9 wash to reduce intestinal bacteria. Worms were centrifuged at 3100 rpm for 2 minutes, then washed with 10 mL of NPB. Worms were then resuspended with prechilled NPB (with 3 μL/mL RNase Inhibitor (Invitrogen) hereafter) to a final volume of 6 mL. All of the Dounce and following steps were performed as for ChIP-seq except for the number of tight strokes (15-22), and two 40 μm cell strainers and two 20 μm cell strainers were used to filter worm debris. Finally, 500 μL TRIzol (Invitrogen) was added to the pelleted nuclei, and then flash frozen in liquid nitrogen and stored at −80°C until RNA isolation.

### Immunostaining

GFP immunostaining was performed on isolated nuclei from OP383, AZ212 and wild type VC2010 worms from six peptone enriched plates for each genotype. Nuclei were isolated as described for ChIP-qPCR and ChIP-seq, except that nuclei were not flash frozen after isolation. One-third of the nuclei was used for an individual immunostaining experiment. All subsequent washes and incubations were performed in 1.5 mL tubes with rotation. 1 mL −20°C methanol was added to isolated nuclei for additional fixation for at least 30 minutes at 4°C. Nuclei were spun at 4000 rpm for 5 minutes and washed 3 times with 1 mL PBST (PBS with 0.1% Tween 20) for 5 minutes per wash at room temperature. Nuclei were blocked with 1 mL 0.5% BSA (American Bioanalytical) in PBST for 30 minutes. Then 120 μL of 1:2000 anti-GFP (ab13970, Abcam) diluted in blocking solution was added and incubated overnight at 4°C. Nuclei were spun at 4000 rpm for 5 minutes at room temperature. Supernatant was removed and 1 mL 0.5 μg/ml DAPI in PBST was added to stain the nuclei for 10 minutes. Nuclei were washed 2 times with PBST for 5 minutes for each wash. 120 μL 1:500 goat-anti-chicken Alex Fluor 488 (A-11039, Invitrogen) secondary antibody in PBST was used to incubate the nuclei overnight at 4°C. Nuclei were washed 3 times with 1 mL PBST for 10 minutes. Nuclei were spun at 4000 rpm for 5 minutes and supernatant was removed. Nuclei were resuspended with 15 μL PBST by gently pipetting. 15 μL antifade mounting medium (Vectashield) was added to the nuclei. 15 μL nuclei were placed on agarose pads under a cover glass with tiny dots of Vaseline on four corners. Images were obtained with a Zeiss Axioplan microscope under an 100X objective and processed with AxioVision software.

### RNA-sequencing

RNA isolation was performed on VC2010 IGN and *glp-1(q224)*, with two biological replicates for each sample. VC2010 germline nuclei were isolated as described for isolation of germline nuclei.

*glp-1(q224)* animals were cultured to starvation on 15-cm NGM plates at 15°C. L1 worms were floated and 10K L1s were plated to one 15-cm NGM plate. Worms were cultured at 15°C for four days until gravid. Adult worms were bleached and embryos were incubated with shaking at 15°C for 36-42 hours. 15K L1s were plated to one peptone enriched plate and cultured at 25°C for 46-48 hours until adult stage. Worms were harvested by washing with M9 3x and were incubated with shaking for 30–45 minutes in M9 buffer. Worms were centifuged at 3100 rpm for 2 minutes. Supernatant was removed and 500 μL TRIzol was added to the worm pellet. Worms were frozen in liquid nitrogen and stored at −80°C until RNA isolation.

Total RNA isolation was performed with standard TRIzol RNA extraction. Approximately 3 μg RNA is typically obtained from each IGN sample collected from six enriched plates. Total RNA was then treated with DNA-*free* rDNase I (Ambion) and cleaned up using RNeasy Mini Kit (Qiagen). rRNA was depleted by Ribo-Zero rRNA Removal Kit (Illumina). The Yale Center for Genome Analysis (YCGA) prepared libraries for each sample using the Kapa Biosystems reagents. At least 20 million 75-bp paired-end reads were acquired for each library using Illumina HiSeq2500.

### RNA-seq analysis

The raw paired-end RNA-seq fastq reads were firstly mapped to rRNA build by bowtie2 (v2.1.0) [37], then the remaining unmapped reads were further aligned to ce10 genome by HISAT2 (v2.0.4) [20] with the mode suppressing the unpaired reads. The gene annotation was downloaded from UCSC Genome Browser, filtered to remove transcripts <50 nt. The expression level of FPKM and significant status were determined by Cuffdiff (v2.2.1) [21]. The bigwig files were generated by SAMtools (v1.3) [38] and BEDtools (v2.17.0) [39], and normalized to 10 million mapped reads for visualization in Genome Browser. All sequencing data are available in Gene Expression Omnibus database under accession GSE117061.

### Preparation of VC2010 IGN and *glp-1(q224)* chromatin and ChIP-sequencing

ChIP-seq of VC2010 IGN was performed using a combined protocol [40, 41] and ChIP-seq on *glp-1(q224)* adult animals was performed as previously reported [41]. VC2010 IGN samples were acquired as described above. 120 μL Nuclear Lysis Buffer (NLB, 50mM Tris pH=8, 10mM EDTA, 1% SDS, 0.5mM PMSF, 2X cOmplete proteinase inhibitor cocktail) was added to each IGN sample. IGN samples were vortexed vigorously for 1 minute and left on ice for 1 minute. The vortex step was repeated. IGN samples were sonicated at 2°C in a water bath sonicator (Misonix S-4000). 20% amplitude and 10 sec on/10 sec off pulses were used for a total processing time of 20 minutes, resulting in enrichment for 100-650 bp DNA fragments. 1.2 mL prechilled FA buffer (50 mM HEPES pH 7.5, 1 mM EDTA, 1% Triton X-100, 0.1% sodium deoxycholate, 150 mM NaCl, add before use: 1 mM DTT, 0.5 mM PMSF, cOmplete proteinase inhibitor cocktail – 1 tablet per 25 mL NPB) was added per IGN sample. 1:20 volume of 20% Sarkosyl solution was added. Sonicated samples were spun at 13,000g for 5 minutes at 4°C. The supernatant was transferred to a new nonstick 1.5 mL tube. 5% of lysate (70 μL) was removed for the input sample and stored at −20°C until the following day to prepare input DNA. 5 μg of anti-H3K27ac (39685, Active Motif) was incubated with each IGN sample overnight at 4°C with rotation.

*glp-1(q224)* animals were cultured as described in RNA-sequencing, except that 50,000 L1s were plated on each peptone enriched plate. Adult *glp-1(q224)* animals from three peptone enriched plates (~1.5×10^5^) were harvested by 3 washes with M9. Worms were crosslinked in 50 mL 2% formaldehyde for 30 minutes in a 50 mL conical tube at room temperature. Formaldehyde was quenched by 1 M Tris (pH 7.5) wash. Worms were then washed 2 more times with M9. Worms were transferred to a 15 mL conical tube and washed with 15 mL prechilled FA buffer. Worms were spun at 3100 rpm for 2 minutes. All but ~200 μL FA buffer was removed and worms were frozen in liquid nitrogen and stored at −80°C. Worm pellets were thawed on ice and 750 μl of FA buffer was added to each sample. Samples were transferred to a 2 mL Kontes Dounce (Kimble Chase). Samples were Dounced 15 times with the small “A” pestle for two cycles with a one minute hold on ice between each cycle. Samples were then Dounced 15 times with the large “B” pestle for four rounds with a one minute hold in between. Samples were transferred to a 15 mL conical tube and FA buffer was added to a final 1.5 mL volume. A quick spin was performed to collect the sample. Samples were sonicated with a SFX250 sonifier (Branson) in an ice bath at 22% amplitude with 10 sec on/1 min off pulses for 34 cycles in a total process time of 5 minutes and 40 seconds. 100-650 bp DNA fragments were enriched after sonication. The sample was transferred to a nonstick 2 mL tube (Ambion) and spun at 13,000g for 15 minutes at 4°C. Supernatant was transferred to a new nonstick 2 mL tube. The protein concentration of the lysate was determined by Bradford assay, and a total of 4.4 mg protein was used for each ChIP sample. Prechilled FA buffer was added to each ChIP sample to bring the volume to 400 μL. 1:20 volume of 20% Sarkosyl solution was added to each ChIP sample. Samples were spun at 13,000g for 5 minutes at 4°C. The supernatant lysate was transferred to a new nonstick 1.5 mL tube. 5% of lysate (20 μL) was removed for input sample and stocked at −20°C overnight. 5 μg of anti-H3K27ac (39685, Active Motif) was incubated with *glp-1(q224)* sample overnight at 4°C with rotation. Both the VC2010 IGN and *glp-1(q224)* samples were treated the same hereafter.

The input samples were thawed the next day and 2 μL 10 mg/mL RNase A (Qiagen) was added to digest the input samples for 2 hours at room temperature. 40 μL (~20 μL of actual beads) protein G Sepharose beads (GE Healthcare) were used for each ChIP sample and washed 4 times with 1mL prechilled FA buffer. The beads were collected with at 2500g for 2 minutes. The entire ChIP sample was transferred to the 1.5 mL tubes with pre-washed beads and rotated at 4°C for 2 hours. Elution buffer (1% SDS in TE, 250 mM NaCl) was added to input samples to bring volume up to 300 μL after RNase A treatment. 2.05 μL of 19.5 mg/mL Proteinase K (Roche) was added to input samples. Input samples were incubated at 55°C for 3 hours. The ChIP samples with beads were washed at room temperature by adding 1 mL of each of the following buffers and incubated for the specified time on a rotator: 2 times FA buffer for 5 minutes; 1 time FA-500mM NaCl (50 mM HEPES pH 7.5, 1 mM EDTA, 1% Triton X-100, 0.1% sodium deoxycholate, 500 mM NaCl) for 10 minutes; 1 time TEL buffer (0.25 M LiCl, 1% NP40, 1% sodium deoxycholate, 1 mM EDTA, 10 mM Tris pH=8.0) for 10 minutes; 2 times TE for 5 minutes. 150 μL elution buffer was added to each ChIP sample and placed in a 65°C heat block for 15 minutes with a brief vortex every 5 minutes. The beads were spun down at 2500g for 2 minutes and the supernatant transferred to new nonstick 1.5 mL tubes. The elution step was repeated and the supernatants were combined. 1.03 μL of 19.5 mg/mL Proteinase K (Roche) was added to each ChIP sample and incubated at 55°C for 1 hour. All input and ChIP samples were transferred to 65°C overnight to reverse crosslinks after Proteinase K treatment. The input and ChIP DNA was purified with a PCR purification kit (Qiagen) following the manufacturer’s protocol. 40 μL TE pH=8 was used to elute DNA.

The Yale Center for Genome Analysis (YCGA) prepared the library and performed sequencing. The KAPA Hyper Library Preparation kit (KAPA Biosystems) was used for ChIP sequencing library prep. DNA fragment ends were repaired with T4 DNA Polymerase, and Polynucleotide Kinase and “A” base added using Klenow fragment in a single reaction followed by ligation of custom adapters (IDT) using T4 ligase. Adaptor-ligated DNA fragments were purified and size selected with Agencourt AMPure XP magnetic beads (Beckman Coulter). Adaptor-ligated DNA fragments were amplified by LM-PCR using custom-made primers (IDT). During LM-PCR, unique 10 base indices were inserted at each DNA fragment and amplified products were purified. The prepped samples were then loaded on to a single-end flow cell and subjected to sequencing. 10-25 million 75-bp single-end reads were acquired for each library using Illumina HiSeq2500 rapid run mode.

### ChIP-seq analysis

The raw ChIP and corresponding input sequencing fastq reads were mapped to WS235 genome by mapping with BWA-map short reads (< 100 bp) (Galaxy Version 0.7.15.1), retaining only high-quality alignments (Q ≥ 20). The bigwig files were generated by bamCoverage (Galaxy Version 2.5.0.0) binned into 25 bp regions and normalized to 1x coverage. Peaks were called by MACS2 (Galaxy Version 2.1.1.20160309.0) with the Minimum FDR q-value=0.01 cutoff for peak detection. The Intersect tool (Galaxy Version 1.0.0) was used to determine the overlapping peaks between IGN-H3K27ac ChIP-seq and SOM-H3K27ac ChIP-seq. Common peaks were defined with an overlap of 1 bp or greater. Metagene analysis custom scripts were used for extracting the value from certain genic regions and averaged for metagene profle by DANPOS [42] with the key parameters (--genomic_sites TSS --flank_up 2000 --flank_dn 2000 --bin_size 50 −exclude P 0.001). Random transcripts were selected for genes with multiple transcription start sites (TSSs). Target calling analysis was performed as described with WS235 annotation [43]. Genes in categories (1,895 ubiquitous genes, 2,230 germline-enriched genes, 169 germline-specific genes, 1,181 soma-specific genes and 858 sperm genes) for metagene profile analysis (Figure 4D) were described previously [13]. Tissue-enriched genes were used to generate a heatmap (Figure 5B) by pheatmap R package.

### RNA-seq and ChIP-seq comparison

ChIP-seq peaks were assigned to target genes as previously described [43]. Several parameters were used to determine the representative peak for target genes that have more than one peak. 1. All peaks with binding at the 3’ end were removed; 2. The peak with the highest ChIP enrichment value was selected if all peaks assigned to the same target were within 1 Kb distance of the TSS; 3. If the highest peak was within 1000-1200 bp, and the ChIP enrichment value is more than 2 times the highest peak within 1 Kb, then it was retained. The unique ChIP peak enrichment value was then determined for each target gene and acquired for correlation analysis with transcript abundance examined by RNA-seq.

### ChIP-qPCR

Samples were prepared as for ChIP-seq, except samples were eluted in 50 uL water. 1 μL of input or ChIP DNA from H3K27ac ChIP experiments, 300 nM of primer, and 12.5 μL FastStart Universal SYBR Green Master (Rox) (Roche) was used in a 25 μL quantitative PCR reaction. ChIP-qPCR were performed as previously described [43]. Relative fold enrichment of germline genes was normalized to negative control *elt-2*. Primers were determined by utilizing ChIP peaks previously identified for L3 N2-H3K27ac ChIP-chip data from the ENCODE project (GSM624432 and GSM624433) [28] or promoter regions with relatively similar distance from the start codon (for negative control).

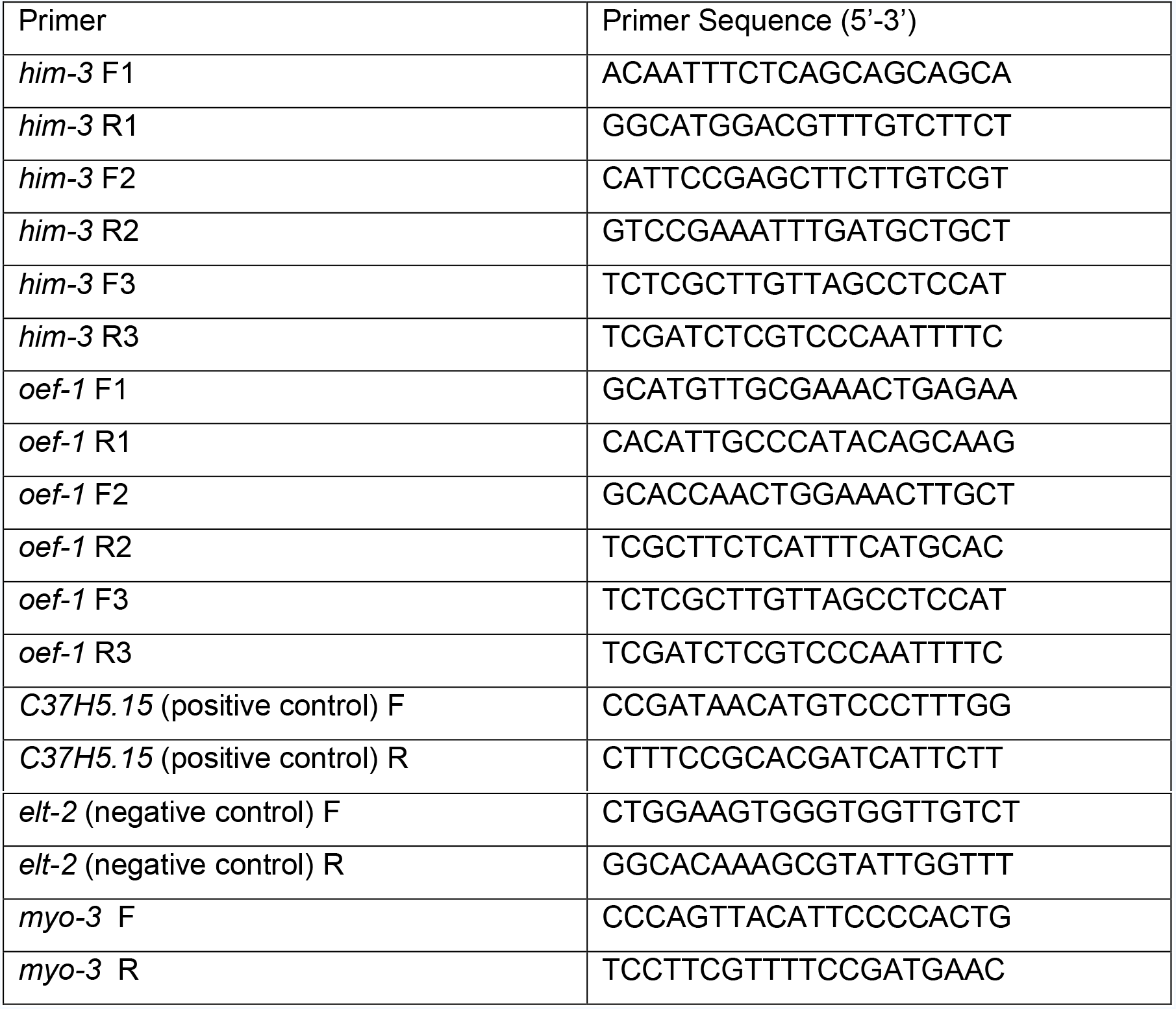
List of ChIP-qPCR primers

## Acknowledgments

We would like to thank James Noonan for equipment use and Guillermina Hill-Teran for technical assistance; The Yale Center for Genome Analysis (YCGA) for RNA-seq and ChIP-seq processing; LaDeana Hillier for target-calling analysis; Robert Waterston, modENCODE and modERN consortium projects and CGC [(NIH) P40 OD-010440] for some strains. This work was supported by NIH R01 GM108663, awarded to V.R.

---

**Figure S1.**
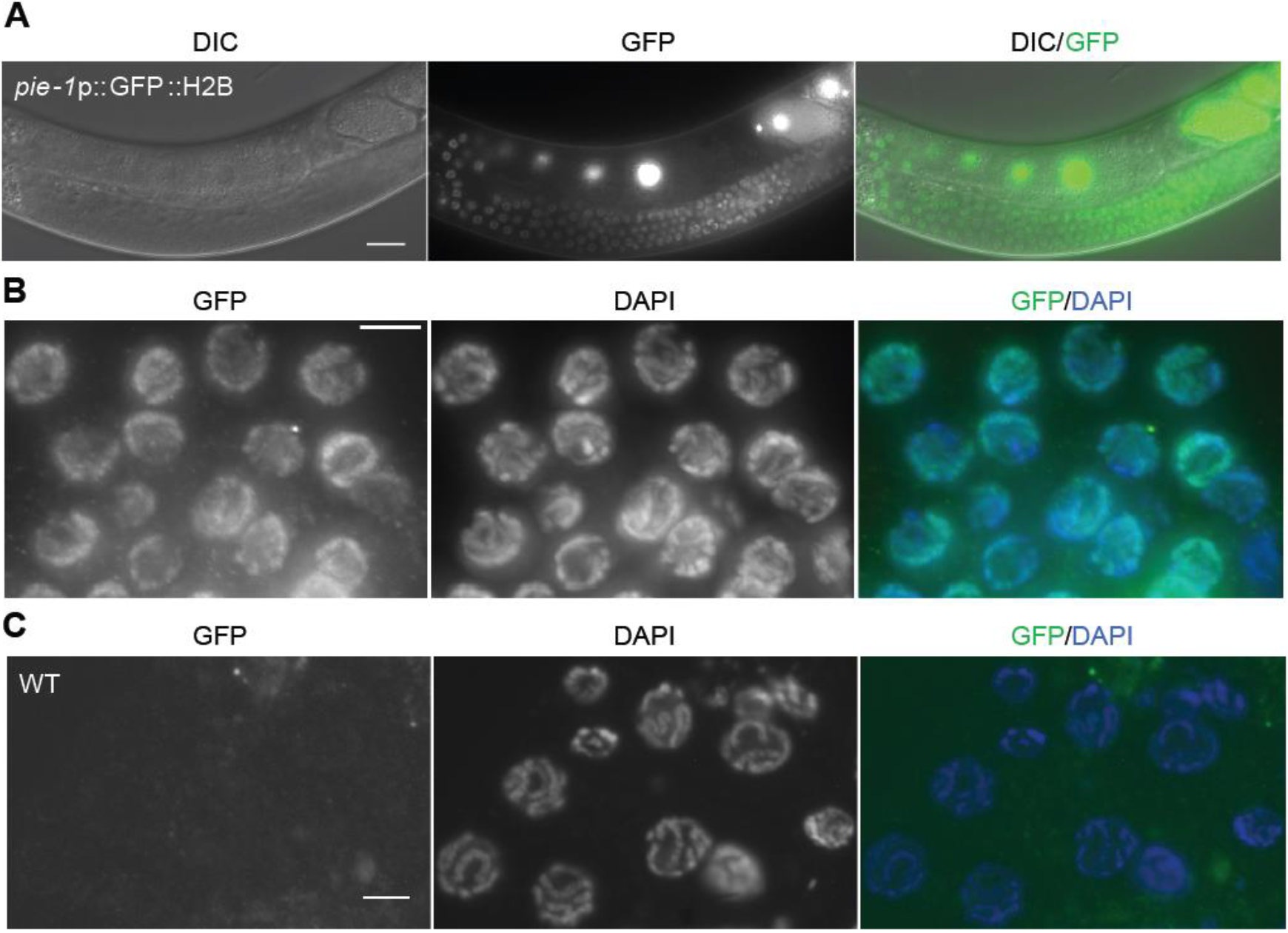
Isolated germline nuclei from a second germline transgenic strain. (A) Young adult *pie-1*p::GFP::H2B transgenic worm showing germline expression. GFP::H2B is expressed in the nuclei of germ cells at all developmental stages, including oocytes and embryos. (B) Isolated nuclei from *pie-1*p::GFP::H2B young adult worms were immunostained with GFP (green) and stained with DAPI (blue). Nuclei stained with both GFP and DAPI (89.1%, n=2111) were considered germline nuclei. Two independent experiments were performed. (C) Isolated nuclei from wild type VC2010 N2 young adult worms were immunostained with GFP (green) and stained with DAPI (Blue). Scale bars, 20 μm in (A), 5 μm in (B) and (C).

**Figure S2.**
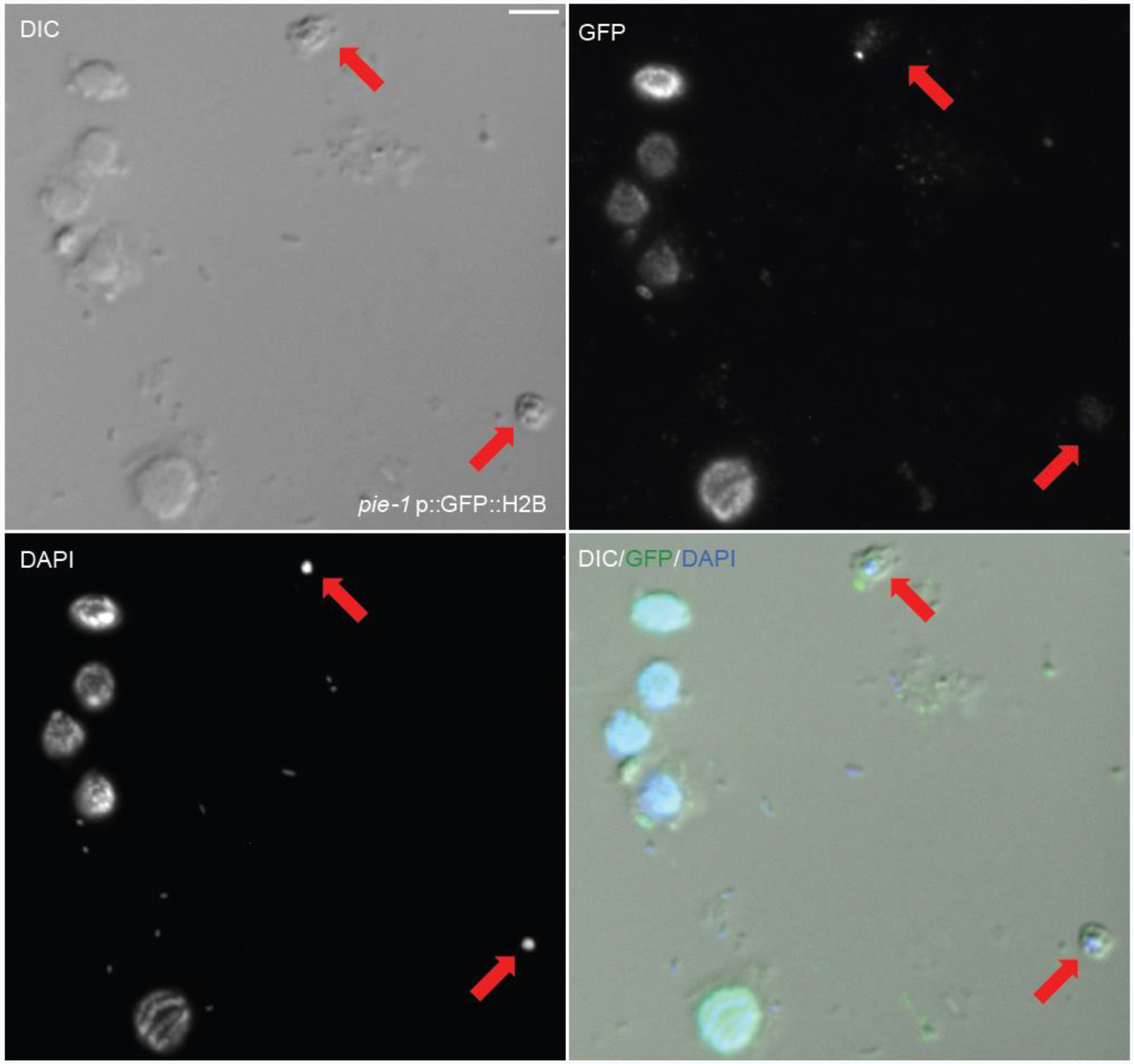
A small fraction of sperm are present in total isolated germline nuclei. Isolated total nuclei from young adult *pie-1*p::GFP::H2B transgenic worms. Nuclei were immunostained with GFP (green) and stained with DAPI (Blue). Arrows indicate two non-GFP stained nuclei with characteristics of sperm. Scale bar, 5 μm.

**Figure S3.**
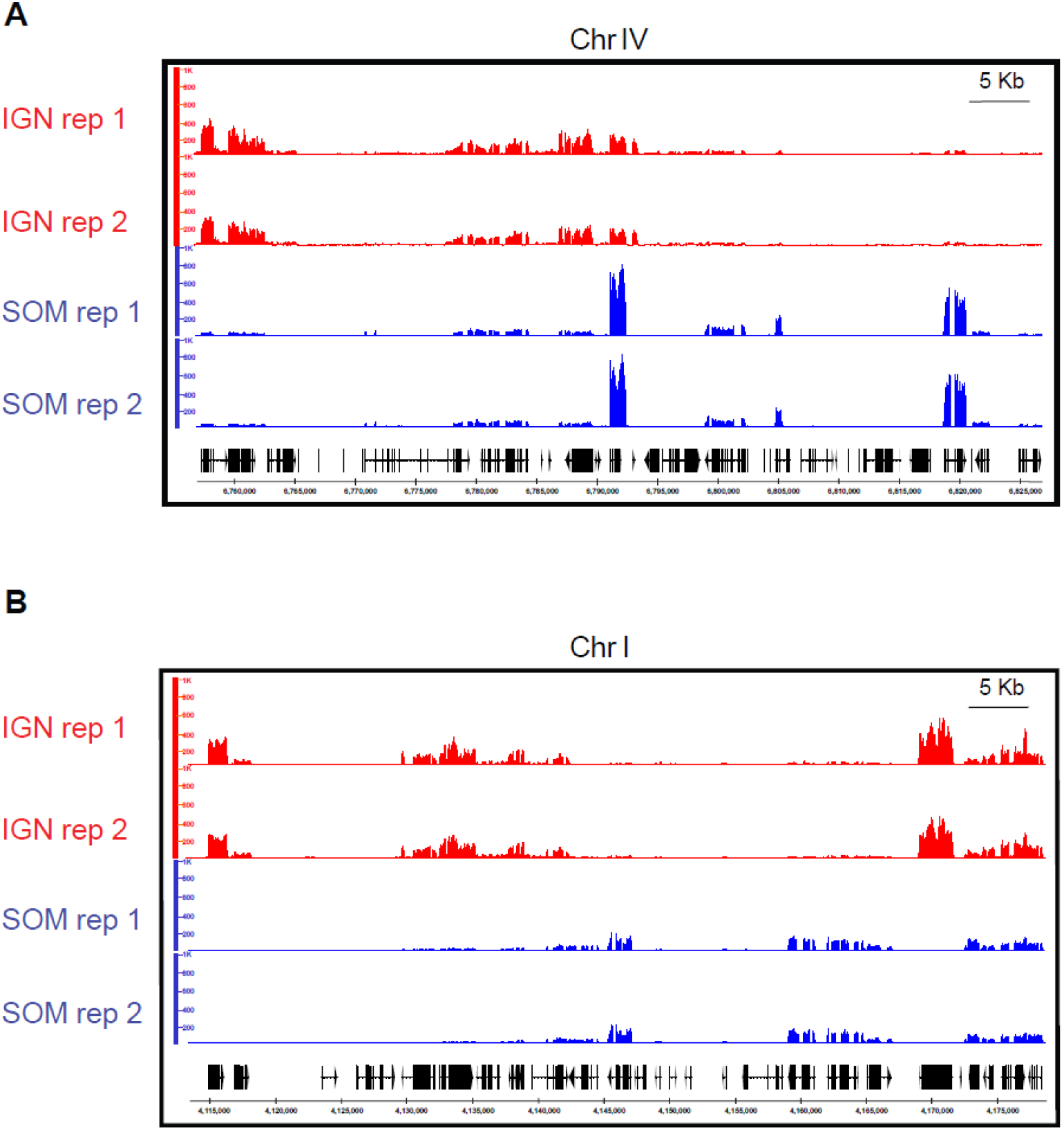
Representative RNA-seq profiles of IGN and SOM. Genome browser views of representative RNA-seq profiles of wild type IGN (red) and SOM (blue) on genomic regions from chromosome IV (A) and I (B), showing tissue-specific transcript abundance.

**Figure S4.**
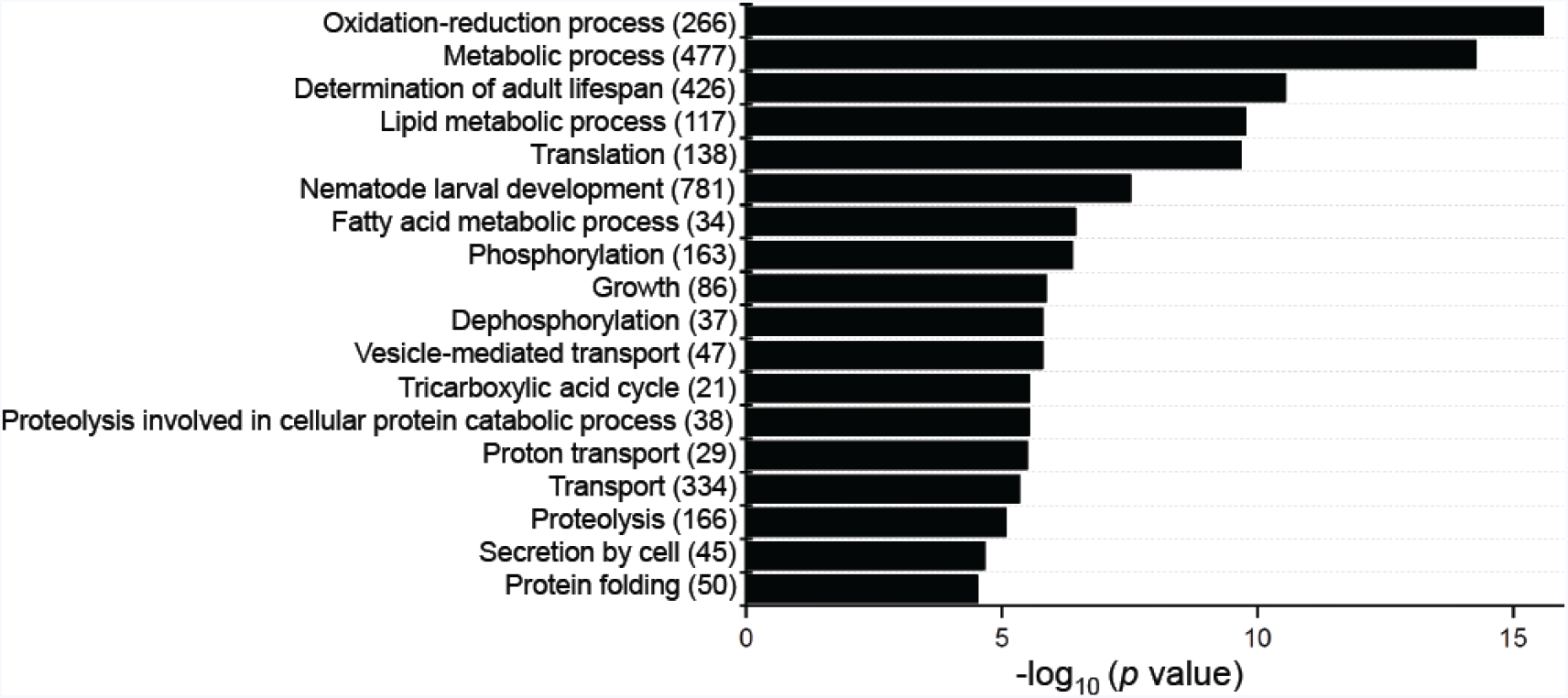
GO analysis of genes present exclusively in the dissected gonad dataset. Eighteen of the most significant Gene Ontology Biological Process terms for the 6606 transcripts present exclusively in the dissected gonad [22] when compared to the IGN dataset in Figure 3D.

**Figure S5.**
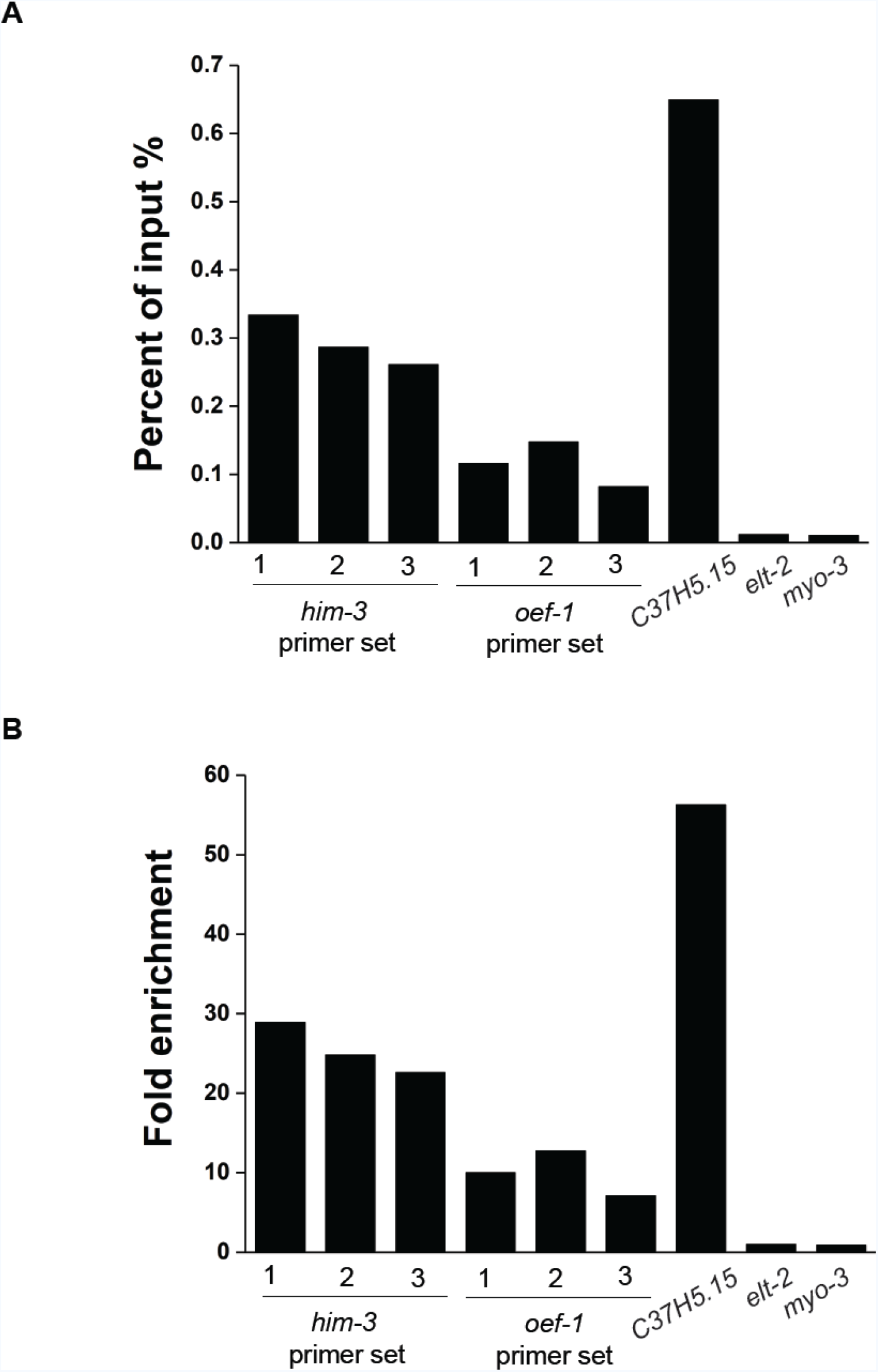
Confirmation of germline-enriched H3K27ac modification in isolated germ nuclei by ChIP-qPCR. Two previously characterized germline-expressed (*him-3* and *oef-1*) and soma-expressed genes (*elt-2* and *myo-3*) were examined for abundance of H3K27ac in IGN by ChIP-qPCR. Three sets of primers were tested for each germline-specific gene. *C37H5.15* served as positive control. *elt-2* served as negative control and was used to calculate fold enrichment. ChIP results are expressed as percent of input using Ct values (A) and fold enrichment of H3K27ac modification normalized to *elt-2* (B).

**Figure S6.**
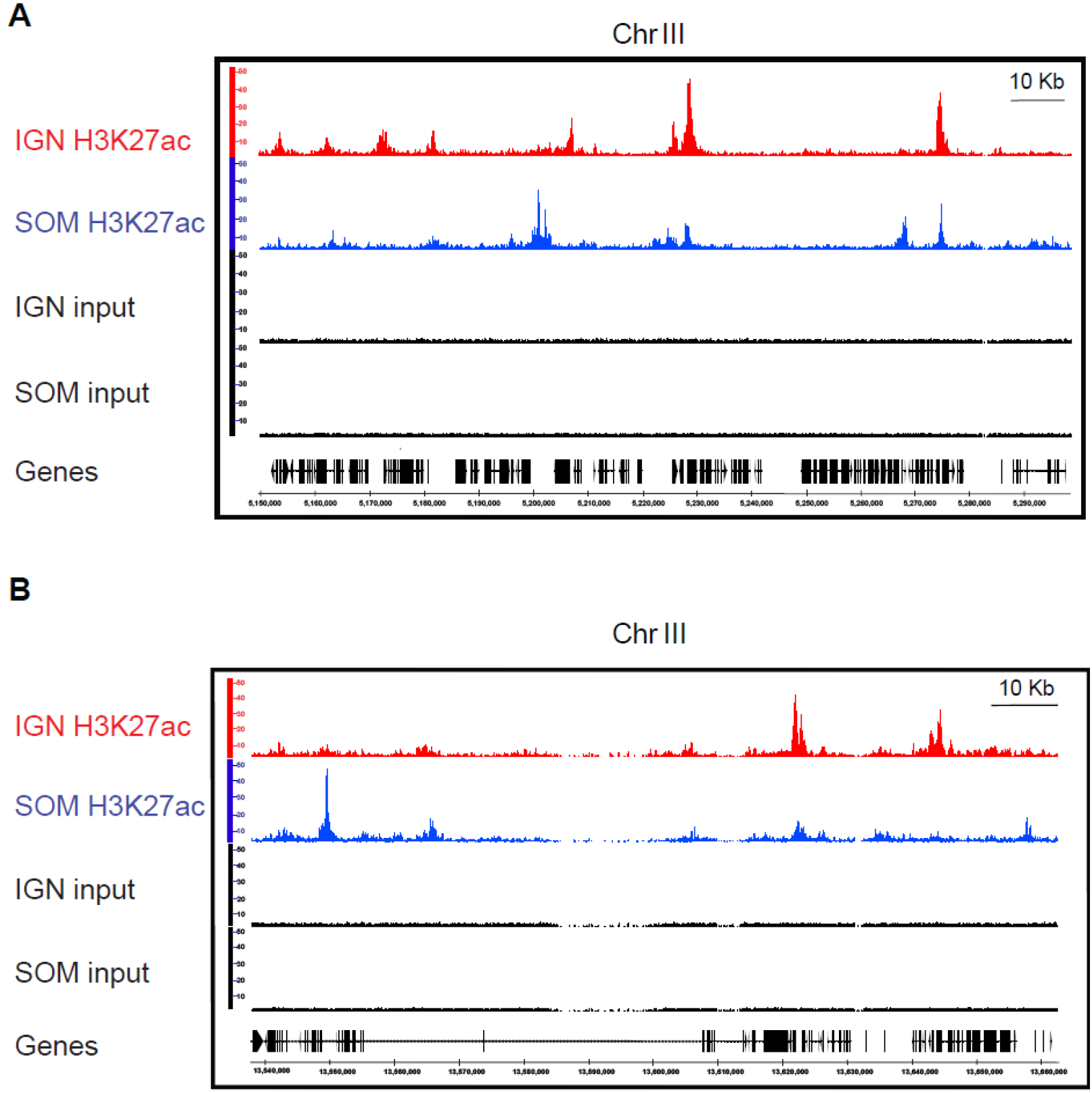
Representative H3K27ac ChIP-seq profiles for IGN and SOM. Genome browser views of H3K27ac ChIP-seq profiling of IGN (red) and SOM (blue) showing representative regions of chromosome III. Input samples are indicated in black.

**Figure S7.**
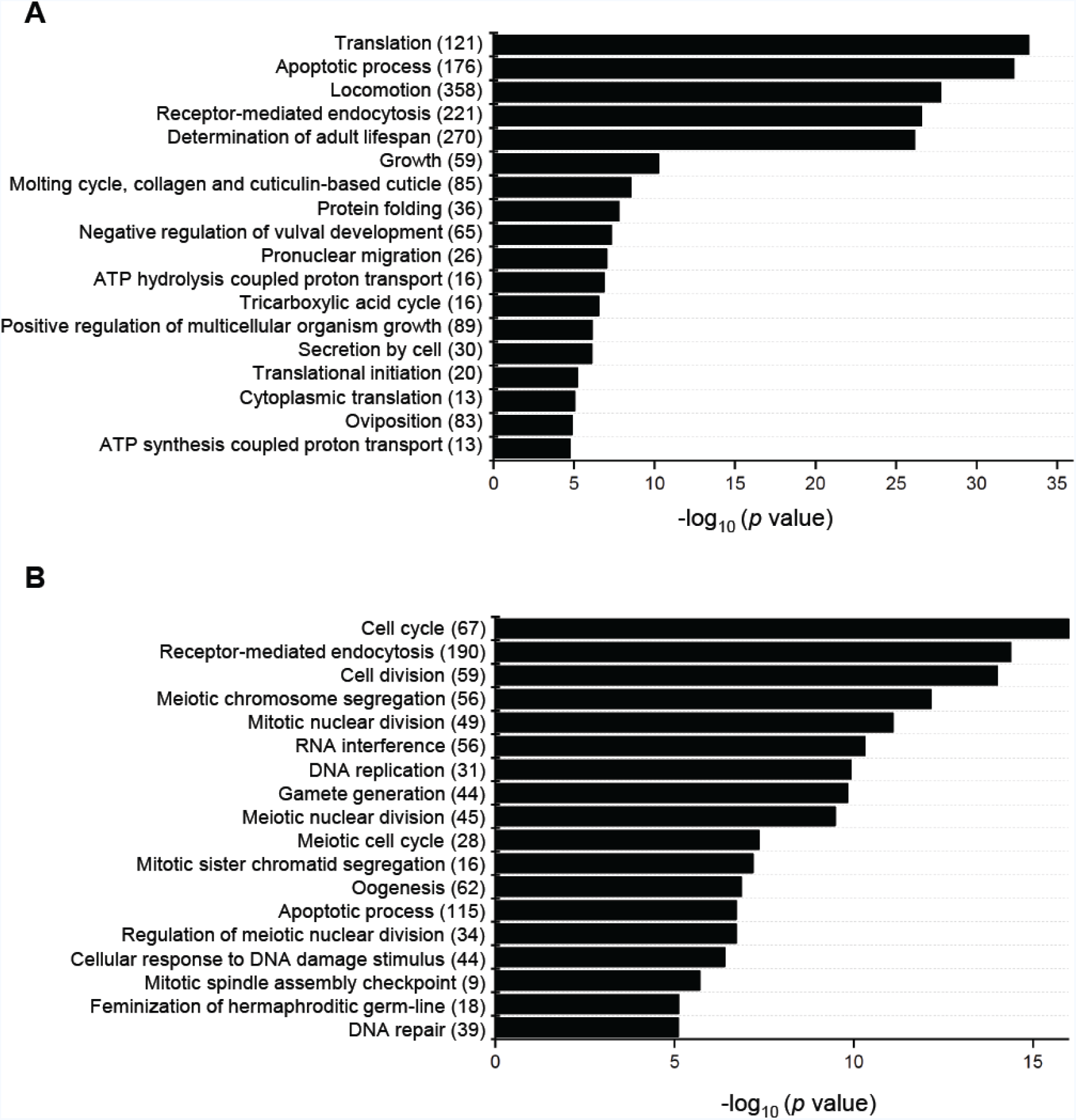
GO analysis of H3K27ac target genes. (A) Eighteen of the most significant Gene Ontology Biological Process terms for H3K27ac target genes with common peaks between IGN and SOM. (B) Eighteen of the most significant Gene Ontology Biological Process terms for IGN-specific H3K27ac target genes.

**Figure S8.**
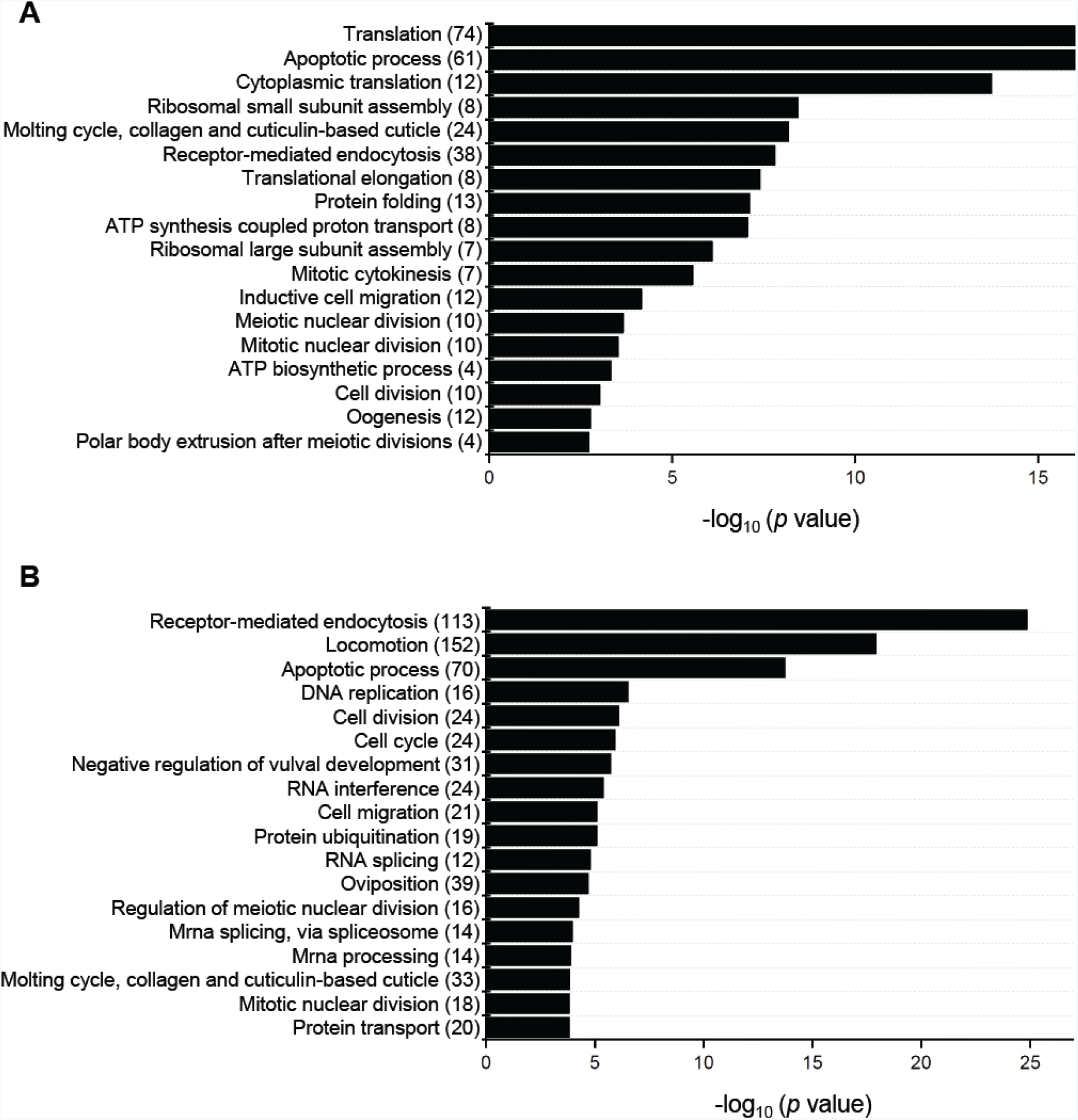
GO analysis of IGN-H3K27ac target genes with high transcript abundance (FPKM≥100) (A) The 18 most significant Gene Ontology Biological Process terms for IGN-H3K27ac target genes with ChIP peak enrichment ≥11. (B) The 18 most significant Gene Ontology Biological Process terms for IGN-H3K27ac target genes with ChIP peak enrichment ≤6.

**Table S1.** Quantification of fraction of germline nuclei. GFP+; DAPI+ nuclei represent germline nuclei and GFP-; DAPI+ nuclei represent non-germline nuclei. Two independent experiments for each genotype were performed.

| Strain | GFP +<br>DAPI + | GFP -<br>DAPI + | Total<br>Nuclei |
| --- | --- | --- | --- |
| OEF-1::GFP | 1936 (91.02%) | 191 (8.98%) | 2127 (100%) |
| *pie-1p::GFP::H2B* | 1881 (89.1%) | 230 (10.9%) | 2111 (100%) |

